# Callose Deficiency Modulates Plasmodesmata frequency and the intercellular space In Rice Anthers

**DOI:** 10.1101/2024.04.08.588242

**Authors:** Harsha Somashekar, Keiko Takanami, Yoselin Benitez-Alfonso, Akane Oishi, Rie Hiratsuka, Ken-Ichi Nonomura

## Abstract

Fertilization relies on pollen mother cells able to transit from mitosis-to-meiosis to supply gametes. This process involves remarkable changes at the molecular, cellular and physiological levels including (but not limited to) remodelling of cell wall. During meiosis onset, cellulose content at the pollen mother cell walls gradually declines with the concurrent deposition of the polysaccharide callose in anther locules. We aim to understand the biological significance of cellulose-to-callose turnover in pollen mother cells walls using electron microscopic analyses of rice flowers. Our observations indicate that in wild type anthers, the mitosis-to-meiosis transition coincides with a gradual reduction in the number of cytoplasmic connections called plasmodesmata. A mutant in the *Oryza sativa* callose synthase GSL5, impaired in callose accumulation in premeiotic and meiotic anthers, displayed a reduction in plasmodesmata frequency among pollen mother cells and tapetal cells suggesting a role for callose in plasmodesmata maintenance. In addition, a significant increase in cell-cell distance between pollen mother cells and impaired premeiotic cell shaping was observed in the mutant. The results suggest that cellulose-to-callose turnover during mitosis-meiosis transition is necessary to maintain cell-to-cell connections and optimal intercellular spacing among anther locular cells explaining the regulatory influence of callose metabolism during meiosis in flowering plants.

**Highlights:**

- Cellulose-to-callose switch during meiosis onset in pollen mother cell walls correlates with changes in cytoplasmic connections called plasmodesmata, among meiocytes and between meiocytes and surrounding somatic cells.
- A mutant in GSL5 (glucan synthase like 5) affects callose deposition but no other cell wall component (e.g., cellulose) in premeiotic and meiotic anthers, except at dyad stage.
- Impaired callose synthesis is proposed to alter plasmodesmata frequency, the extracellular spacing and the shaping of pollen mother cells during mitosis-to-meiosis transition.
- Increase in plasmodesmata frequency negatively correlates with cell-cell distance between pollen mother cells in both wildtype and a callose mutant.

## Introduction

During the reproductive phase of flowering plants, spore mother cells abort somatic division to enter into meiosis: a specialized cell division stage through which chromosomes are halved and haploid spores are produced. The mitosis-meiosis transition determines the steady supply of gametes for fertilization, making it pivotal for plant reproduction. This transition involves an extensive remodelling of pollen mother cells (PMCs, the male meiocytes) within the anthers (male reproductive organs), such as cell size enlargement, adoption of a spherical morphology, and replacement of cellulose-rich walls for callose (a *β*-1,3 glucan polysaccharide) (Helpson-Harrison, 1964&1966; Kelliher & Walbot, 2011; Matsuo et al. 2013; Unal et al. 2013). Another noticeable event during this transition is the cell-cycle synchronization, wherein the asynchronous mitotic division cycle among sporogenous cells (i.e., PMC founder cells), is turned to a synchronous meiotic cycle within the anther locule (Whelan, 1974). However, the relationship between the mitosis-meiosis transition and the drastic changes associated with PMC shape and properties during this cell cycle switch remains poorly understood.

In rice anthers, callose is synthesized at the PMCs-facing locular centre by a membrane-spanning callose synthase, the *Oryza sativa* GLUCAN SYNTHASE LIKE 5 (OsGSL5) and accumulates at the intercellular spaces among PMCs and the surrounding somatic tapetal cells (TCs), shielding PMCs from the somatic cell niche (Somashekar et al. 2023). The marked accumulation of callose is a prominent histological hallmark of PMCs exiting mitosis and entering meiosis in plants. Though the significance of callose for pollen formation and development has been well documented (Albert et al. 2011; Franklin-Tong, 1999; Prieu et al. 2017; Qin P et al. 2012; Zhang et al. 2018), its biological significance in male meiosis initiation and progression was only recently uncovered (Somashekar et al. 2023; reviewed in Somashekar & Nonomura, 2023). The *Osgsl5* mutant anthers lack callose, leading to a precocious initiation of premeiotic DNA replication, followed by several meiotic defects in chromosome condensation and behaviour, homolog synapses and pollen development (Shi et al. 2015; Somashekar et al. 2023; Somashekar & Nonomura, 2023). These defects result in complete male sterility, suggesting the essential roles of callose for mitosis-meiosis transition and pollen formation.

Callose is a glucan polymer joined by *β*-1,3 glycosidic bonds and sporadic *β*-1,6 side branches (Stone and Clarke, 1992; Chen and Kim, 2009; Zavaliev et al. 2011; Piršelová and Matušíková, 2013; Nedhuka, 2015). It is deposited in the newly formed cell plate during cell division (Staehelin and Heplar, 1996; Hong et al. 2001; Thiele et al. 2009), in sieve plates controlling sieve pore size in phloem tissue (Xie et al. 2011), as part of the mechanisms for plant defence against invading pathogens and abiotic stresses (Dong et al. 2008; Voigt, 2014; Jacobs et al. 2003), during pollen tube elongation (Adhikari et al. 2020), among other processes. Callose is also an integral part of plasmodesmata (PDs) which are membranous channels/bridges connecting the cytoplasm of neighbouring cells in most plant organs/ tissues, forming the symplasmic pathway for molecular transport (Barnett, 1982; Beebe et al. 1992). Callose regulates PDs conductivity, and thereby, the transport of molecular signals between cells (Radford et al. 1998; Lucas et al. 2009; Lee and Sieburth, 2010; Zavaliev et al. 2011; Sager and Lee, 2018). Callose is also proposed to regulate the physicomechanical properties of cell walls thus affecting the diffusion of molecules across the extracellular (apoplastic) space (Maltby et al. 1979; Bhalla and Slattery, 1984; Yim and Bradford, 1998; Abou-Saleh et al. 2018).

In this study, we aim to understand the biological significance of callose regulation in PMCs during the mitosis-meiosis cell-cycle switch in rice anthers. We used electron-microscopy to visualize PMC-PMC, PMC-TC and TC-TC boundaries in anther locules. We found abundant PDs connecting these cell interfaces in premeiotic anthers filled with callose walls in locules. During transition to meiosis, the PD frequency (number of PDs per cell wall area) was gradually reduced at PMC-PMC interfaces, while remained comparable at PMC-TC interfaces between premeiotic and meiotic anthers. In comparison to wildtype (WT), the callose-synthesis mutant *Osgsl5* displayed a reduced PD frequency connecting PMCs and surrounding TCs in both premeiotic and meiotic anthers. In addition, we observed an increase in the intercellular space between PMCs within the anther locules in the *Osgsl5* mutant when compared to WT. Our results indicate that the level of callose deposition in the PMC walls during mitosis-meiosis transition impacts PD formation or maintenance and influence the apoplastic space (and the shape) in the meiocytes, likely affecting the diffusion of molecular signals required for proper male meiosis initiation and progression. We discuss how this mechanism might be linked to the defects in male meiosis and pollen development observed in the rice *Osgsl5* mutant and propose new avenues to investigate the role of PDs in synchronizing mitosis-meiosis transition, gamete formation and, thereby, plant fertility.

## Material and Methods

### Plant materials and growth conditions

The *Osgsl5-2* and *Osgsl5-3* biallelic mutants previously produced by CRISPR-Cas9-mediated target mutagenesis of *OsGSL5* gene (Os06g0182300) (Somashekar et al. 2023) were used in this study. The *japonica* rice cultivar “Nipponbare”, from which the *Osgsl5-2* and *Osgsl5-3* mutants originated, was used as the wild type. Plant materials were grown at 30°C day and 25°C night temperature and 70% relative humidity with 12 hr daylength in growth chambers.

### Aniline blue staining

Rice florets were fixed in 4% (w/v) paraformaldehyde (PFA)/1× PBS. The lemma and palea of florets were removed with anthers kept still intact with the floret base. Anthers were then dehydrated in graded ethanol series (30%, 50%, 70%, 90% and 100%) for 30–60 min each. Samples were then embedded with Technovit 7100 (Kulzer Technique) according to the manufacture’s protocol. The hardened embedded sample sections of thickness 4-6µm were taken using the microtome R2255 (Leica). Sections air dried and stained with 0.01% (v/v) aniline blue (Sigma Aldrich) in 0.1 M K_3_PO_4_(pH 12) for 25–30 min. Sections were observed under a confocal laser scanning microscope system (FV300; Olympus) for callose, and images were processed with ImageJ (https://imagej.nih.gov/ij/) (Schneider et al. 2012). Callose immunostaining in premeiotic anther sections with monoclonal anti-β-1,3-glucan (callose) antibody (Biosupplies Australia) performed as described by Somashekar et al. (2023). A dilution of 1/1000 for primary antibody and 1/200 for CY3-conjugated anti-mouse IgG (Merck) labelled secondary antibody was used for staining.

### Cellulose staining

For observation of cellulose accumulation pattern in anthers at different meiosis stages, WT, *Osgsl5-2* and *Osgsl5-3* flowers were fixed and processed as above. Technovit embedded samples provided for plastic sectioning in 4-6µm thickness using the ultramicrotome RM2255. The sections were air dried and stained with Renaissance stain 2200 (Renaissance chemicals Ltd, UK) which highlights cellulosic cell walls. Fluorescent images captured by the confocal laser scanning microscopy Fluoview FV300 (Olympus) were processed with ImageJ software (https://imagej.nih.gov/ij/) (Schneider et al. 2012).

### Transmission Electron Microscopy

Young rice flowers were prefixed in 4% paraformaldehyde (PFA) in 1x PBS buffer (pH 6.8-7.0) followed by a second fixation in 1% PFA and 2.5% glutaraldehyde (GA) in 0.05M PBS (pH 6.8-7.0) at 4 °C overnight. After washing with 0.05M PBS, flowers were further fixed in 2% (w/v) osmium tetraoxide (OsO4) in 0.05M PBS for 1hour on ice, and then washed with 0.05M PBS (pH 6.8-7.0). Dehydration was done in a graded ethanol series of 50%, 70%, 80% and 90%, for 30 minutes each, followed by overnight shaking in 90% EtOH. Next day the sample was further dehydrated in 95% and twice in absolute EtOH for 30 minutes each. The dehydrated sample was transferred to 1:1 propylene oxide (PO) and absolute EtOH for 30 min and then immersed twice in pure PO for 30 min. Samples were incubated with Quetol 812 (Nisshin EM) and PO in ratio of 1:1 for 120 min and 2:1 overnight, followed by incubation in the pure resin at 37°C for 120 min and further in fresh pure resin at 37 °C for 60 min. The sample was embedded in a flat embedding mold with the pure epoxy resin, and incubated at 60°C for 2 days. The resin block was trimmed, and ultrathin sections of 100-150 nm were made by the ultramicrotome EM UC6 (Leica). Sections were first stained with 4% w/v uranyl acetate for 10 min, and with lead stain solution (Sigma-Aldrich) for 5 min. Sections were observed under transmission electron microscopy (TEM) JEM1010 (JEOL).

### Quantification of PD frequency on Electron Microscopy images

PD numbers at PMC-PMC, PMC-TC and TC-TC interfaces were quantified in two different anther stages roughly corresponding to premeiotic interphase (Meiosis stage 1 (Mei1) = 0.35-0.45mm anther length) and early meiotic stages (Meiosis stage 2 (Mei2) = 0.50-0.55mm anther length). The PD frequency was calculated by dividing PD numbers against a given cell-cell distance (measured in µm). Statistical significance for PD frequency at each cellular interface between WT and *Osgsl5-3* anthers was calculated by Student’s t-test. At least 3 independent rice florets were observed for each stage in WT and *Osgsl5-3* anthers.

### Measuring cell-to-cell distances on electron microscopic images and quantifying callose signal intensity

Distance between two adjacent cells at three cellular interfaces viz., PMC-PMC, PMC-TC, and TC-TC, was obtained by measuring the cell-cell distance between two adjacent cell membranes at a given cellular interface. The average distance at all three cellular interfaces in WT and *Osgsl5-3* was taken. For normalization, the absolute cell-cell distance values of three interfaces were divided by the average of TC-TC distances measured at the same meiotic stage for the same genotype, because a difference in TC-TC distances was largely comparable between Mei1 and Mei2 stages and between WT and *Osgsl5-3* anthers. The statistical significance of normalized cell-to-cell distances was estimated by Mann Whitney’s U test.

### Quantification of callose and cell-cell distance in anther sections

The callose immunostaining and TEM images obtained by the methods above mentioned were used for measurement of fluorescence signal intensity and cell-cell distance at early premeiotic interphase stage. In premeiotic anthers which contain PMCs mostly in angular shape, a PMC-PMC interface was divided into 13 bins from the central side (locule center/central side) to the lateral TC side. The PMC-PMC distance at each bin was measured using ImageJ. For quantification of the callose signal, the line was drawn along the apoplastic space of the PMC-PMC interface, where callose is deposited, from the central side to TC side by the ImageJ option “segmented line (thickness 5)”, and the pixel intensity (intensity profile) was obtained alongside the line by the multi plot option of ImageJ. Pixel values were normalized by the average of all pixel values measured at the same PMC-PMC interface and averaged for each bin.

## Results

### Callose deposition during mitosis-to-meiosis transition is disturbed in *Osgsl5-2* and *Osgsl5-3* mutant anthers

Callose deposition is a histological hallmark of pollen mother cells initiating meiosis in flowering plants. To monitor the callose deposition during mitosis-to-meiosis transition phase, we performed aniline blue staining. Callose was absent in wild type (WT) mitotic anthers – Mitosis stage (Mit = <0.35mm anther length) (Fig. 1A-C) but it is found highly deposited among PMC cells and between PMC and surrounding tapetal cells during premeiotic interphase stage – Meiosis stage 1 (Mei1 = 0.35-0.45mm anther length), thus shielding the germ cells completely from the outer somatic cell niche (Fig. 1D). Two anther callose defective mutants in rice – *Osgsl5-2* and *Osgsl5-3* were previously characterized (Somashekar et al. 2023). As previously reported, both *Osgsl5-2* and *Osgsl5-3* displayed almost complete loss of callose around PMCs, and between PMCs and surrounding tapetal cell layer (Fig. 1E-F). During early meiotic prophase I stage – Meiosis stage 2 (Mei2 = 0.50-0.55mm in length), callose can be seen accumulating at the center of anther locule where PMCs face each other in WT, whereas, in both *Osgsl5-2* and *Osgsl5-3* anthers, callose is severely disturbed with very faint to almost no signal of callose observed at locule center (Fig. 1G-I). This observation supports the role of GSL5 as the main regulator of callose biosynthesis during cellulose-to-cellulose turnover during mitosis-meiosis transition in rice anthers.

**Figure 1:**
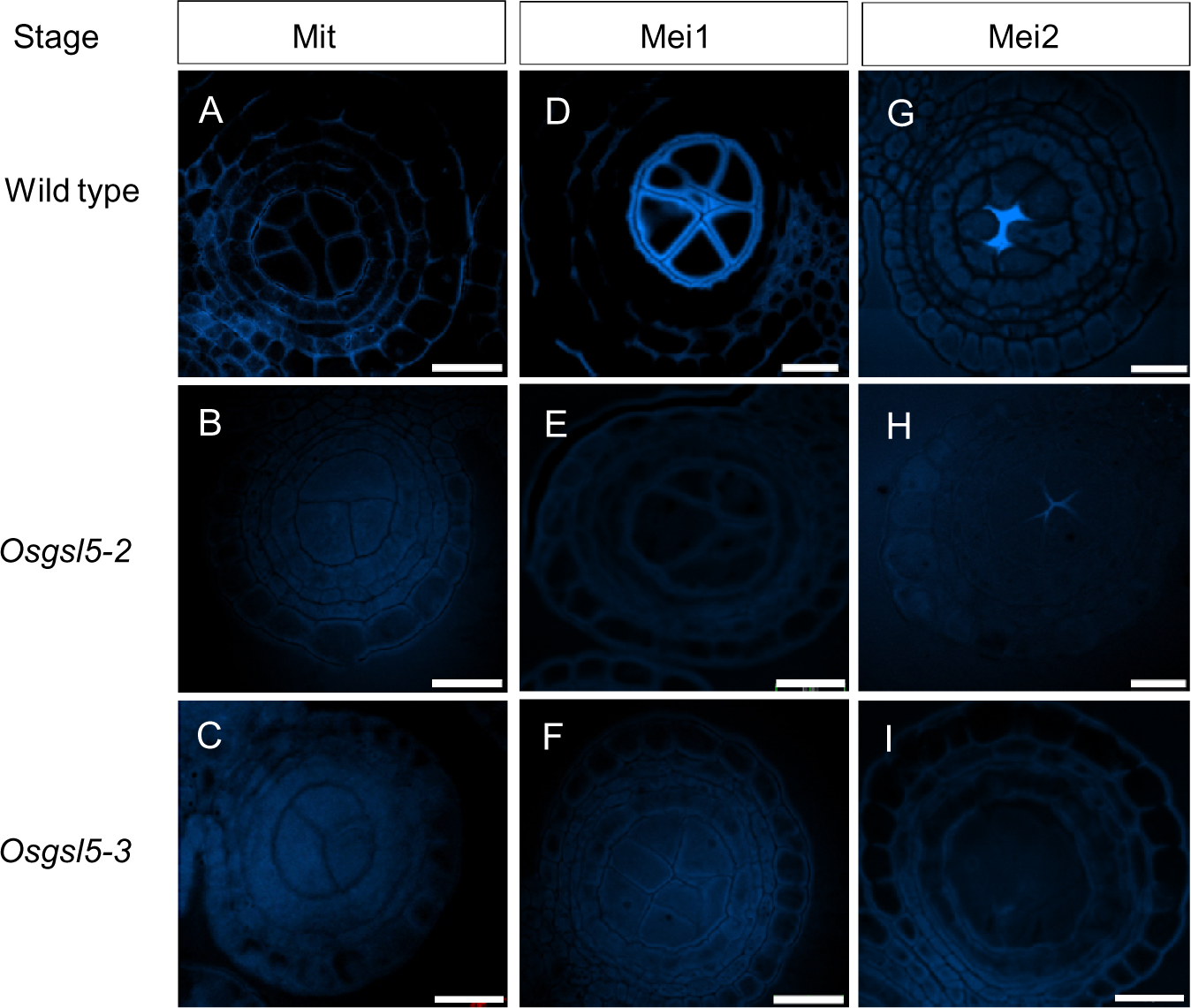
Callose deposition during mitosis-meiosis transition in wild type (WT), *Osgsl5-2* and *Osgsl5-3* in rice anthers. During mitotic sporogenous cell stage (Mit, anther length (AL) =< 0.35mm), callose is not present within the locular space of anthers in WT, *Osgsl5-2* and *Osgsl5-3* (A,D,G). At premeiotic interphase stage (Mei1, AL= 0.35-0.45mm), callose is deposited around the pollen mother cells in WT anther locules but not in *Osgsl5-2* and *Osgsl5-3* anthers (B,E,H). During early meiotic prophase stages (lepto/zygo/pachy, Mei2, AL= 0.5-0.55mm), the callose is deposited at the center of locule in WT anther (C,F,I). Very little to no callose is observed at early prophase stage in *Osgsl5-2* and *Osgsl5-3.* Scale bar=20µm.

### PD frequency varies across three cellular interfaces during mitosis-meiosis transition

The importance of intercellular communication during mitosis-meiosis transition and between two consecutive stages of meiosis (Mei1 and Mei2) is unknown. To investigate extracellular structures involved in communication, we carried out TEM experiments in anther locules at three cellular interfaces viz., PMC-PMC, PMC-TC and TC-TC (Fig. 2) in two different anther stages Mei1 (0.35-0.45mm in length) and Mei2 (0.50-0.55mm in length) roughly corresponding to premeiotic interphase and subsequent early meiosis I stages respectively, as previously defined (Itoh et al. 2005). A previous study evidenced that the anther length-based temporal expression of two TC-specific genes, *TIP2* and *EAT1*, and the layered structure of premeiotic and meiotic anthers were unaffected by the *Osgsl5* mutation (Somashekar et al. 2023), thus anther length was selected as a good standard parameter to compare premeiotic interphase and meiotic events between WT and *Osgsl5* anthers. We observed a number of PDs as electron-dense tubular structures bridging neighbouring locular cells and calculated PD frequency as a parameter to quantify these phenotypic observations. At the PMC-PMC interface, the PD frequency was relatively high during Mei1 stage (0.112/µm in average), but it was gradually reduced as PMCs transit to meiosis during Mei2 (0.072/µm in average). PD frequency remained largely unchanged at PMC-TC interface (0.104 and 0.103/µm in average) in both Mei1 and Mei2 stages (Fig. 2). On the other hand, PD frequency at TC-TC interface increased two-fold in Mei2 anthers (1.030/µm in average) when compared to Mei1 anthers (0.517/µm in average) (Fig. 2).

**Figure 2:**
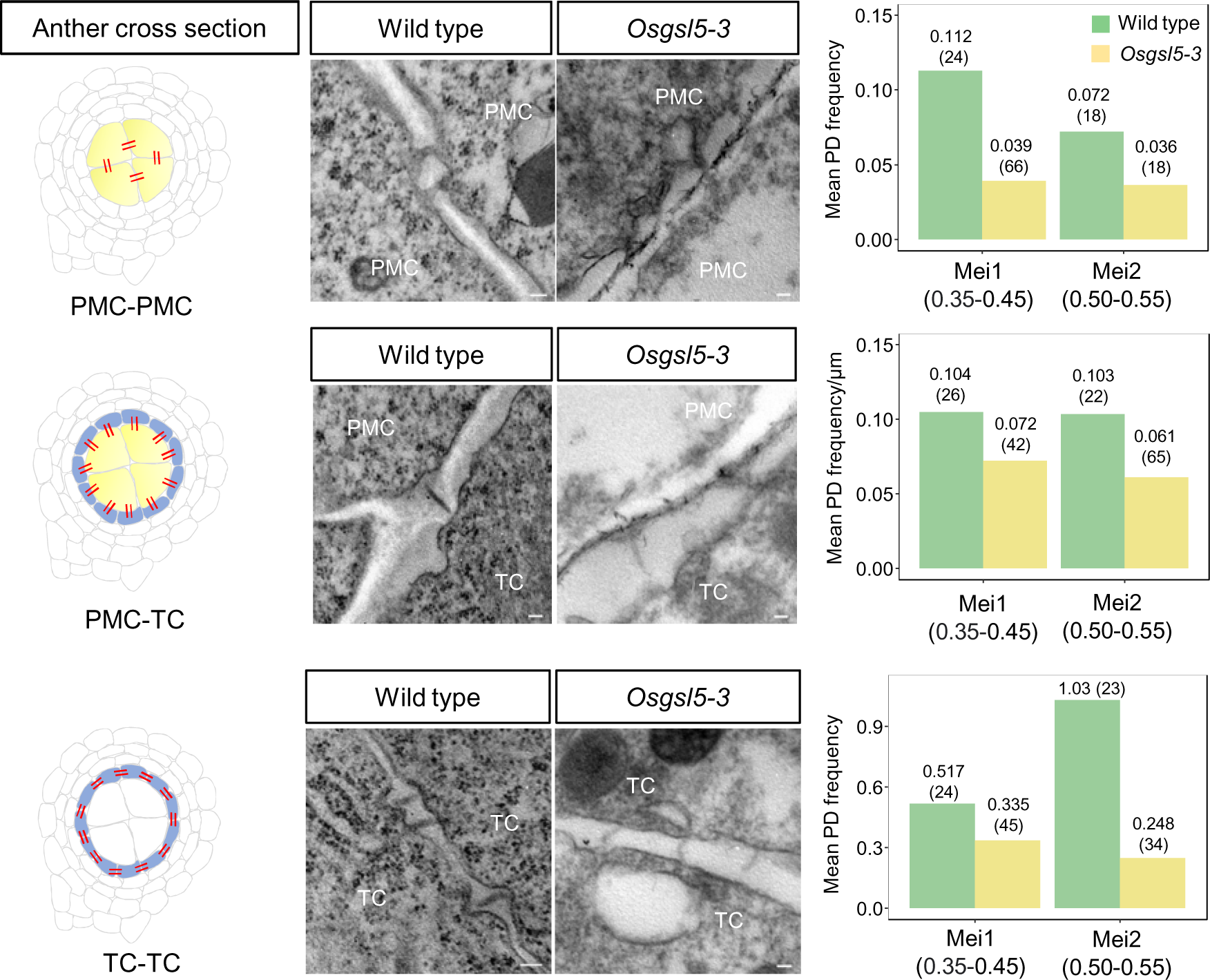
Plasmodesmata frequency is differentially regulated upon entry to meiosis and in *Osgsl5-3* mutant anthers. The left cartoons depict cross view of premeiotic anthers (the double lines in red indicate plasmodesmata, PD) showing the different cellular interfaces between pollen mother cells (PMC) and tapetum cells (TC). Middle panels: transmission electron microscopy images showing PDs at the three cellular interfaces in wild type and *Osgsl5-3* anthers. Right panels: quantification of PD frequency in Mei1 (premeiotic interphase) and Mei2 (early meiosis I) for wild type and *Osgsl5-3* anthers. Mean PD frequency (#/μm of cell wall) are shown on top of the bars and in bracket indicate the number of sections quantified for PD frequency at respective anther stage in wild type and *Osgsl5-3* anthers. Anther lengths (mm) used for Mei1&2 are indicated in the brackets below the labels. ML: middle lamella. Scale bar= 100nm.

In our pictures, the membrane at the vicinity of PDs looked invaginated towards the cytoplasmic side (Fig. 2). Such membrane invaginations were more often observed at PDs located at the TC-TC interface than at the PMC-TC and PMC-PMC interfaces. At PMC-PMC and PMC-TC interfaces, PDs were mostly single or rarely double, whereas PDs at TC-TC interface quite often occurred in clusters (Fig. 1, Fig. S1). No PDs were observed in any of the studied interfaces at later meiosis stages (0.7mm in anther length) presumably because they are severed (Fig. S2).

In addition, qualitative comparison of the size (aperture) of PDs connecting PMCs, found that these are larger compared with PDs bridging PMC-TC and TC-TC interfaces and, in some instances, diffusion of what appear cellular organelles was observed among the PMCs in premeiotic anthers (Fig. S3).

In summary, our observations in WT anthers, suggest a downward trend of PD density at the PMC-PMC interface as PMCs transit from mitosis to meiosis, but not at the PMC-TC or TC-TC interfaces. At the TC-TC interface, in fact, the PD density shows an upward trend (Fig. 2).

### A rice mutant in callose synthesis shows reduced PD frequency in anthers during mitosis-meiosis transition

Callose is believed to be involved in the regulation and maintenance of PDs occurring between PMCs in anthers (Albertsen & Palmer, 1979; Echlin & Godwin, 1968; Mamun et al. 2005; Steer, 1977; Sager & Lee 2014). Therefore, we asked whether callose deposition in premeiotic interphase anthers influence the distribution of PDs. For TEM analyses, we used the rice mutant *Osgsl5-3* severely affected in callose synthesis (Fig. 1F), and strongly defective in meiosis initiation (Somashekar et al. 2023, Supplemental material). We calculated PD frequency in anthers at Mei1 and Mei2 stages from TEM images.

In *Osgsl5-3* mutant anthers, the PD density was, in general, reduced when compared to WT at all three cellular interfaces and at both Mei1 and Mei2 stages (Fig. 2) although statistically the difference was non-significative (p=0.92, 0.32, 0.23 at PMC-PMC, PMC-TC, TC-TC respectively). Unlike WT, PDs in *Osgsl5-3* remained at comparable density between Mei1 and Mei2 (0.039/µm and 0.036/µm in average at PMC-PMC, 0.072/µm and 0.061/µm in average at PMC-TC, and 0.335/µm and 0.248/µm in average at TC-TC, respectively) (Fig. 2). Noteworthily, while in WT the PD density at PMC-PMC and PMC-TC interfaces was similar, in *Osgsl5-3* anthers the density was less than half at the PMC-PMC than at PMC-TC interfaces at both Mei1 and Mei2 stages.

### Cell-Cell distance among PMCs increased in the callose synthase mutant *Osgsl5-3*

When analysing TEM pictures in *Osgsl5-3* anthers, we often observed a dramatic increase of intercellular spacing at PMC-PMC interface in comparison to WT anthers (Fig. 3A). We measured the cell-cell distance at the three cellular interfaces in WT and *Osgsl5-3* anthers to test a hypothetic role for callose synthesis in apoplast spacing.

**Figure 3:**
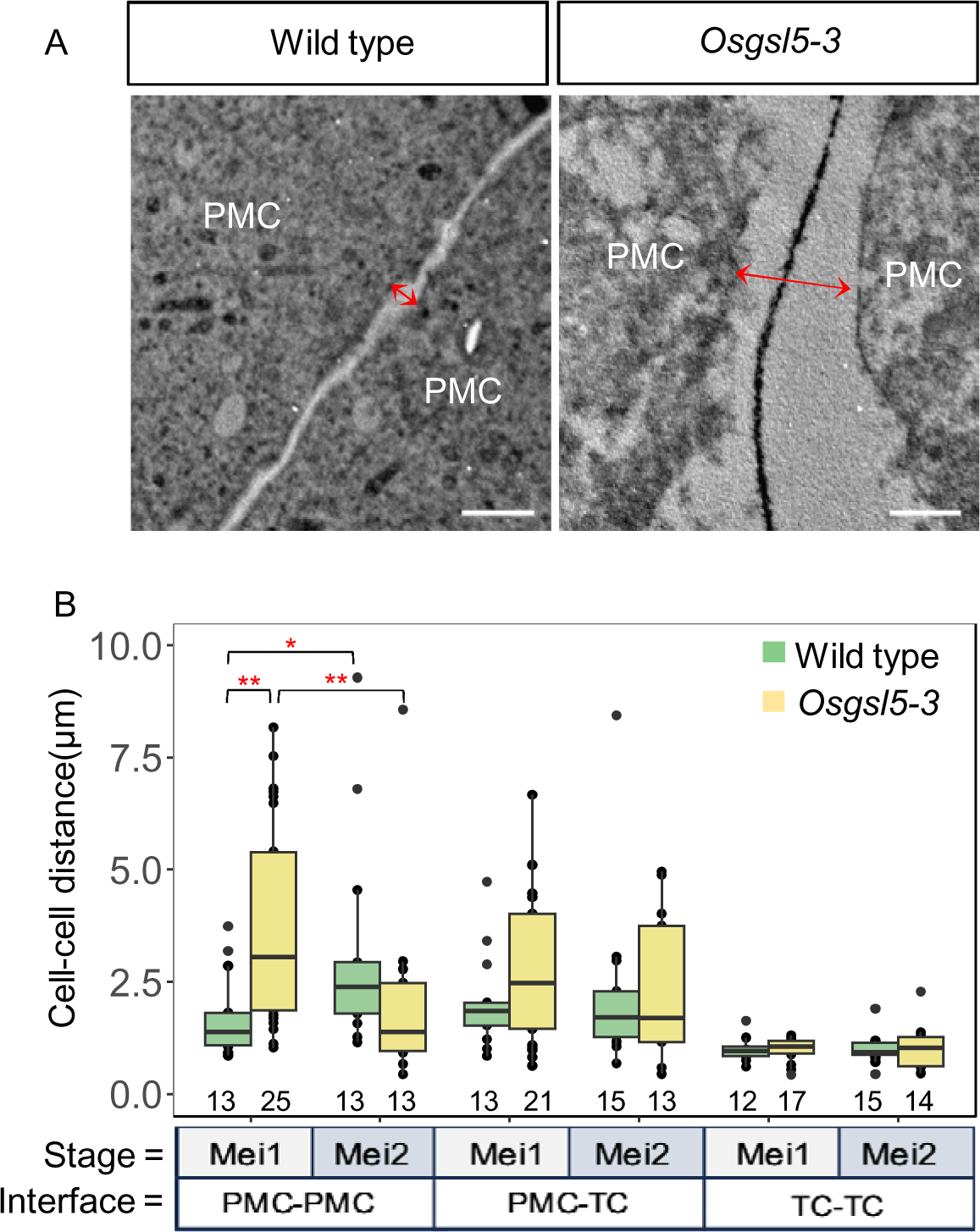
Extracellular distance between pollen mother cells increases upon entry into meiosis in Wild type (WT) and in *Osgsl5-3* anthers. A: Representative electron micrographs showing the cell-cell distance (red double arrowhead) between two adjacent PMCs in WT and *Osgsl5-3* anthers. Dark gray line between cells is the pectin enriched middle lamella (ML). Scale bar = 1µm. B: Comparison of cell-cell distance (µm) at three cellular interfaces between WT and *Osgsl5-3* anthers. All cell–cell distances are normalized against TC-TC distance. Horizontal line within each box indicates the average cell-cell distance. Asterisks indicate significant differences (\**p* ≤ 0.05, \*\**p* ≤ 0.01, \*\*\**p* ≤ 0.001 by Mann Whitney’s U Test). Premeiotic interphase (Mei1) anther = 0.35-0.45mm in length, early meiosis I (Mei2) anther = 0.50-0.55mm. Numbers below the plot correspond to the number of sections observed for each interface and stage in both WT and *Osgsl5-3*.

As PMCs progress from Mei1 to Mei2, the angular PMCs move apart from each other and become spherical, resulting in increased apoplastic spaces among WT locular cells. Our measurements indicate a significant increase in the apoplast space at PMC-PMC interface in WT Mei2 anthers (3.14µm in average, p=0.03) compared to Mei1 anthers (1.75µm in average). In contrast, the extracellular space at both the PMC-TC and TC-TC interfaces remained almost unchanged (2.07 µm and 2.25 µm in average at PMC-TC interfaces in Mei1 and Mei2, respectively and 1µm in average at TC-TC in both Mei1 and Mei2) (Fig. 3B).

In *Osgsl5-3* Mei1 anthers, the increment in apoplastic space between PMCs was significantly higher than WT at the PMC-PMC interface (3.60µm in average, p=0.002) (Fig. 3B). A similar trend was observed at PMC-TC interface in Mei1 anthers, but the difference was non-significative (2.81µm in average, p=0.22). In *Osgsl5-3* Mei2 anthers, the PMC-PMC distance (2.06µm) was reduced compared to WT Mei2 (3.14µm) and significantly reduced compared to *Osgsl5-3* Mei1 anthers (3.60µm, p= 0.008) (Fig. 3B). The TC-TC distance remained mostly comparable through both stages in both WT and mutant anthers (Fig. 3B).

Both callose and cellulose are synthesized from UDP-glucose, thus a change in cellulose-to-callose turnover might be due to an increase in cellulose (associated with more substrate available due to defective callose biosynthesis). To determine if cellulose accumulation contributes to the increase in the apoplastic spaces, we used renaissance staining (which stain cellulose in cell walls). No major differences were observed between WT and both *Osgsl5* mutant anthers during mitosis-meiosis transition, except at dyad stage (Fig. 4). Similar to the WT, in both *Osgsl5-2* and *Osgsl5-3* anthers, cellulose rich cell walls are observed at PMC and somatic cells during mitosis-meiosis transition (Fig. 4A, E&I). During early meiosis stages, cellulose disappears from PMCs but remain present in soma cell layers (Fig. 4B, C, F, G, J&K). Intriguingly, during the dyad stage, the newly formed cell plates showed reduced cellulose in the *Osgsl5* mutant anthers compared to WT (Fig. 4D, H&L).

**Figure 4.**
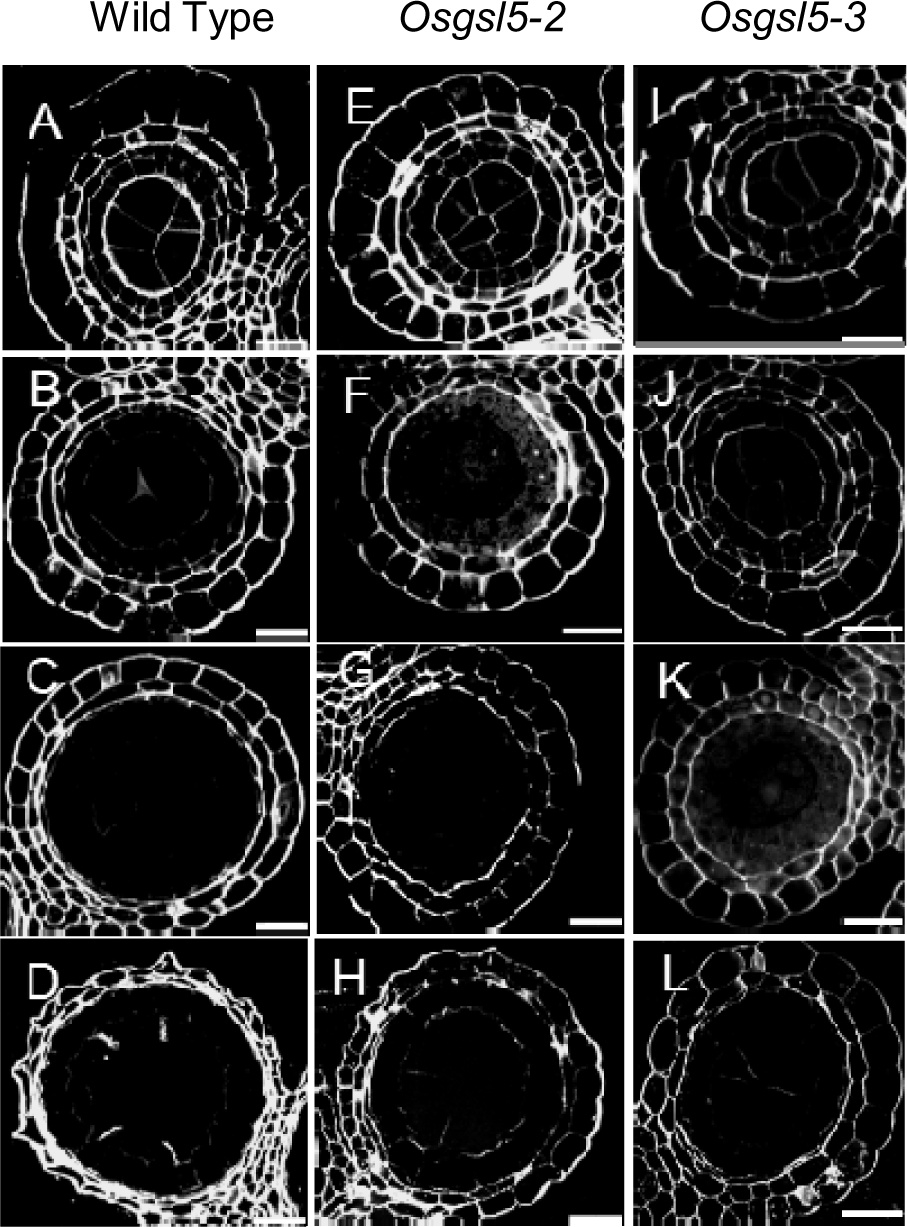
Cellulose accumulation in Wild type (WT), *Osgsl5-2* and *Osgsl5-3* anthers. Renaissance staining revealed cellulose fluorescence in WT, *Osgsl5-2* and Os*gsl5-3* cell walls of pollen mother cells undergoing mitosis (A, E, I), premeiosis interphase (B, F, J), Early-mid prophase I (C, G, K) and at the dyad stage (D, H, L). Scale bar = 20µm.

Together these observations indicate that depletion of callose in the mutant premeiotic anthers influences (directly or indirectly) the apoplastic space at PMC-PMC and PMC-TC interfaces and the effect is stronger at the PMC-PMC interface than at the PMC-TC interface (consistent with regions where OsGSL5-dependent callose accumulates (Somashekar et al. 2023). At these early stages of meiosis, cellulose content does not appear affected suggesting that mutation in callose synthesis does not severely influence general cell wall composition until the dyad stage.

### PD frequency negatively correlates with cell-cell distance among anther locular cells

Depletion of callose in the *Osgsl5* mutant anthers were linked to reduced PD densities and enhanced cell-cell distance. To investigate the relationship between callose, PD number and cell-cell distance, we identified correlations between these parameters at all three interfaces in WT and *Osgsl5-3* anther locules. Scatter plots in Fig. 5 display PD density as a function of PD frequency in each cell-cell interface. Roughly PD frequency negatively correlated with cell-cell distance at PMC-PMC (*r*=-0.289 and −0.313 in WT and *Osgsl5-3* respectively) and PMC-TC interfaces (*r*=-0.207 and −0.388 in WT and *Osgsl5-3* respectively) while at the TC-TC interface there is no obvious correlations (Fig. 5). This effect appears independent of callose. Despite the obvious reduction in PD frequency discussed above, correlation coefficients in *Osgsl5* mutant were similar to WT.

**Figure 5:**
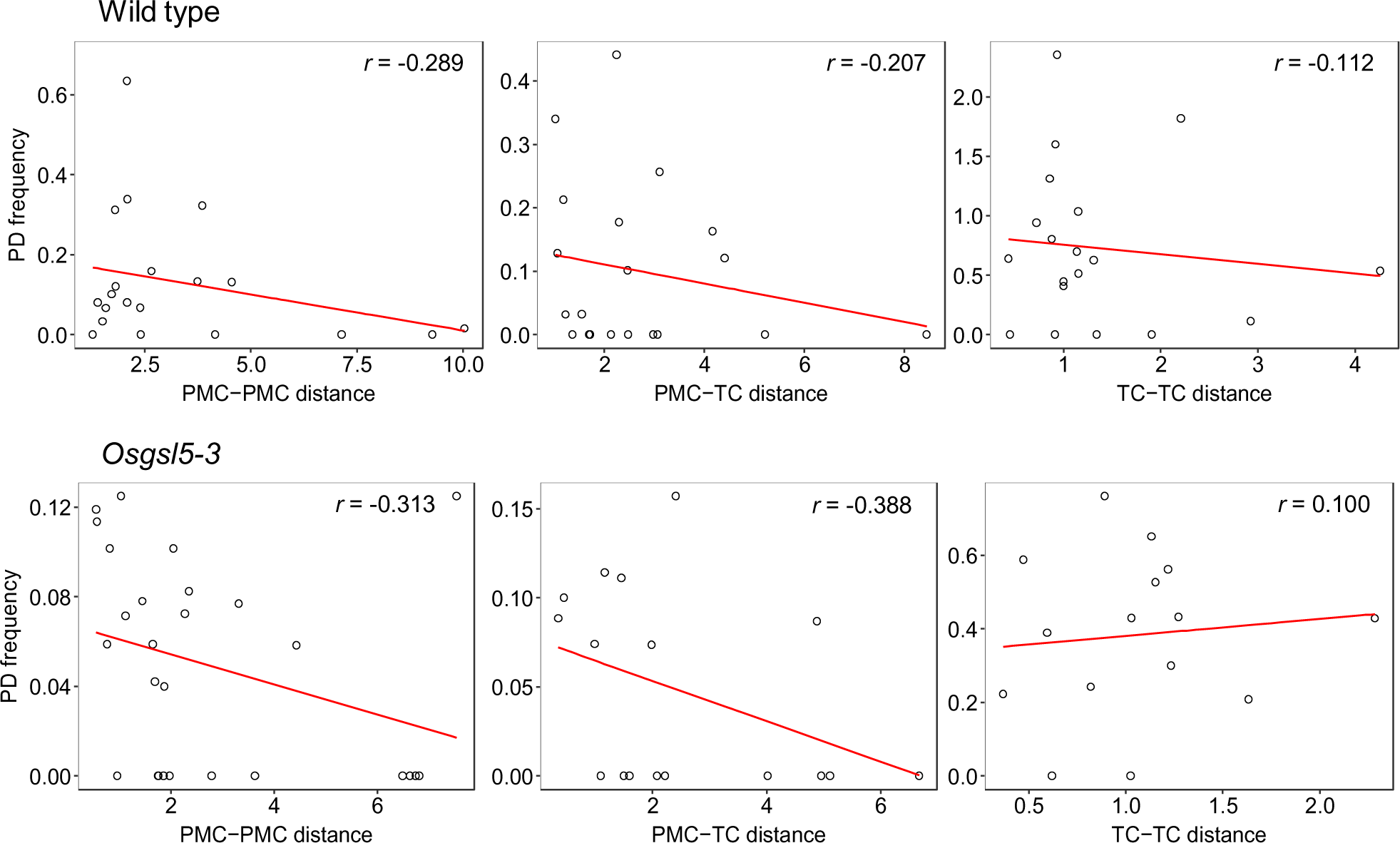
Correlation between cell-cell distance and PD frequency at three cellular interfaces in Wild Type (WT) and *Osgsl5-3* anthers. Scatter plot showing relationship between PMC-PMC (left), PMC-TC (middle), TC-TC (right) distances and PD frequency (#/µm cell wall length) in WT (above) and *Osgsl5-3* (below) anthers respectively. The PD frequency is plotted against the given cell-cell distance (µm) at each interface. The correlation coefficient values (r) for each combination of cell-cell distance and PD frequency are shown on the plots. The data from both premeiotic interphase and early meiotic-I anthers is used for correlation analyses between PD frequency and cell-cell distance.

These observations suggest a potential link between the presence (or dissolution) of PDs and the physical cell-cell distances among PMCs and between PMCs and TCs.

### Mutation in callose synthesis modulates the shaping of pollen mother cells during the mitosis-meiosis transition

In WT premeiotic interphase stage (0.45mm anthers), we often found an increased PMC-PMC distance at the intercellular space proximal to the locular center (central side) while the distance at the distal side near TCs remained narrow (Fig. 6A&B). This stage corresponded to anthers undergoing a beginning stage of callose deposition (according to determinations reported in Somashekar et al. 2023 Supplemental material). When a corner of PMCs starts to curve at the proximal region, another corner at the distal side maintained an angular shape and this region of apoplastic enlargement accumulates callose detected using immunolocalization (Fig. 6A). To further investigate the role of callose synthase in initiating the widening of the PMC-PMC distance at the locular center, we divided a PMC-PMC interface into thirteen bins, and in each bin, we quantified the cell-cell distance and the callose immunofluorescence intensities along the interface from proximal (the locular center/central side) to distal regions (the TC side) (see Methods section). We found that the PMC-PMC distance largely corresponds to the sites of callose deposition along the PMC-PMC interface (Fig. 6A’, B). Conversely, when cell-cell distance was calculated in the *Osgsl5* mutant across the same interface (locular center to TC side), no major changes were observed (Fig. 6C). in WT, we observed the initial sites of callose deposition at the locule center, not only on the cross sections but also on the longitudinal sections of the anther (Fig. S4). Callose deposition was often greater at the longitudinally central side of PMC-PMC junction than that at the apical or basal side in anther locules, suggesting a high polarity of PMCs with respect to callose deposition on both anticlinal and periclinal directions.

**Figure 6:**
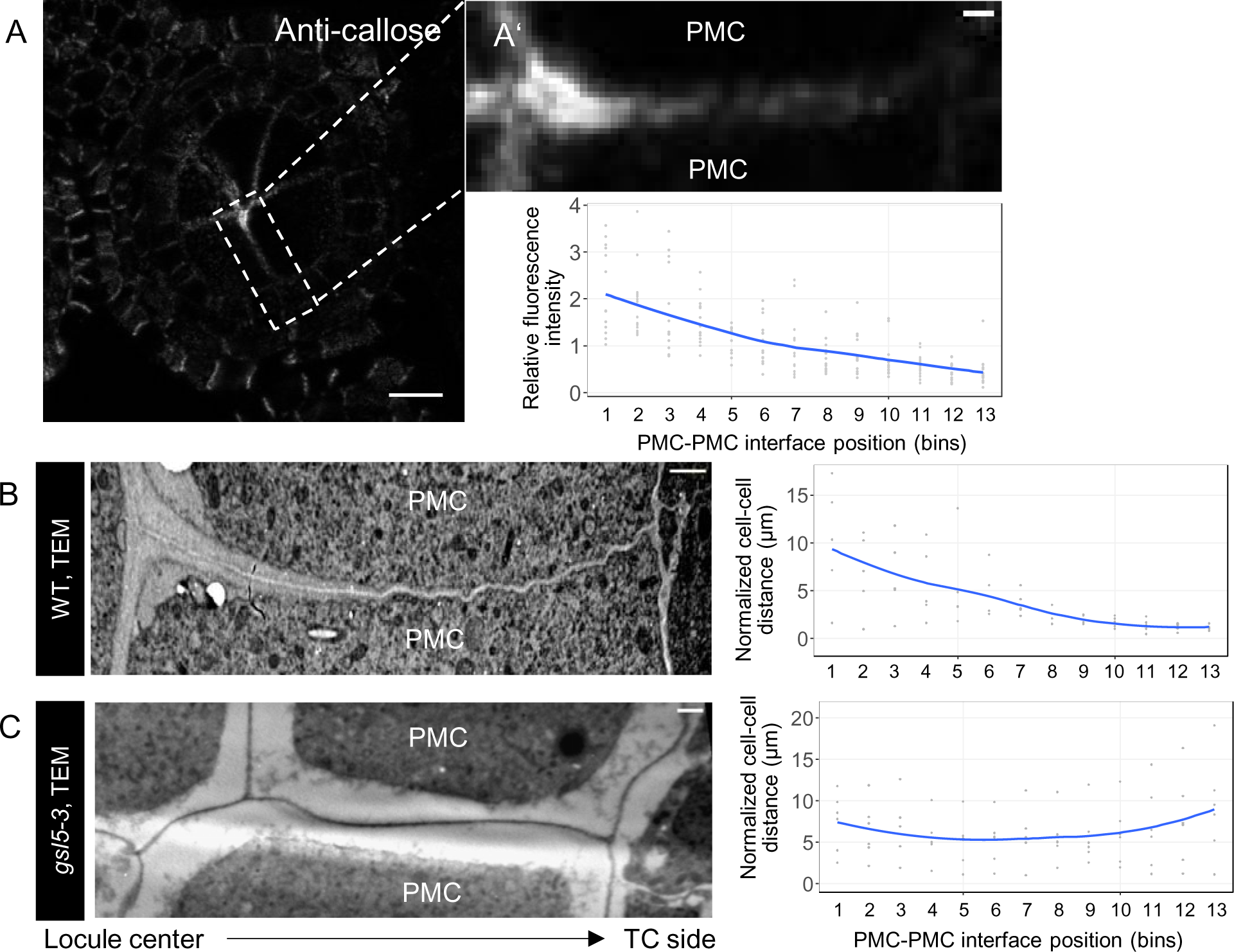
Callose deposition and cell-cell distance in premeiotic Wild type (WT) and *Osgsl5-3* mutant anthers underpin differences in the shaping of pollen mother cells (PMC). A. Image showcasing the callose deposition at the PMC-PMC junction in WT early premeiotic interphase anthers prior to spreading to the PMC-TC interface. Callose is immuno-stained using anti-callose antibody. On the right (A’), the picture shows a zoom in to highlight the differential spreading of callose in the PM-PMC interface. On the bottom graph, fluorescence intensity (as a measure of callose deposition) is quantified along this interface. Below left are TEM section of early premeiotic WT anthers (B) and *Osgsl5-3* mutant anthers (C). The graphs on the right show the PMC-PMC cell-cell distances across the interfaces. Note the downward curve associated with PMC shaping in WT while cell-cell distances remained large but constant across the PMC-PMC interface in the mutant. Scale bar (A) =10µm, (A’, B, C) = 1µm.

The results suggest that, besides affecting PD number and cell-cell distance, callose uneven distribution might play a role in the shaping of PMCs in early premeiotic anthers.

## Discussion

### Cellulose-to-callose switch in pollen mother cells at the onset of meiosis is necessary to regulate PD frequency

In this study we aimed to understand the importance of cellulose-to-callose switch in PMC walls during the mitosis-to-meiosis transition in rice anther locules. Using electron-microscopy, our observations revealed relatively abundant PDs at the premeiotic interphase in PMCs, corresponding to the stage where callose accumulation occurs within WT anthers (Mei1) (Fig. 1&2). In early meiotic prophase I anthers (Mei2), the PDs gradually declines, thus reduction in PDs correlate with callose deposition upon entry into meiosis (Fig. 1&2). This is supported by past studies reporting that the dissolution of cytoplasmic connections between PMCs is likely due to callose deposition (Albertsen & Palmer, 1979; Echlin & Godwin, 1968; Mamun et al. 2005; Steer, 1977; Sager and Lee, 2014). In contrast, the interface between TCs displayed completely the opposite trend with Mei2 anthers showing higher PDs than Mei1, suggesting an increase in symplastic connectivity between TCs after meiosis initiates (Fig. 2). One previous study claimed no observable PDs in PMC-TC interface in rice premeiotic anthers (Mamun et al. 2005). However, we found PDs at PMC-TC interfaces (at similar frequency as found between PMCs) in premeiotic and meiotic anthers (Fig. 2), consistent to observations made in anthers of other plants like oats, pepper and plum meiocytes (Horner and Rogers, 1974; Radice et al. 2008, Steer,1977). Our results suggest that all PMC-PMC, PMC-TC and TC-TC interfaces are symplastically connected but PD formation (or dissolution) is differentially regulated during mitosis-meiosis transition in each of these interfaces. Maintenance of PDs could be necessary to allow symplastic flow of information from surrounding somatic cells to the meiocytes (Heslop-Harrison, 1964 & 1966; Heslop-Harrison& Mackenzie, 1967; Plackett et al. 2014; Liu et al. 2017, Zhai et al. 2015) and/or, the other way around, from sexual cells to somatic cell layers (Yang et al. 1999). Although the nature of the symplastically mobile molecules has not been yet identified, PDs are pathway for transcription factors, hormones and RNAs (including silencing signals) that regulate important developmental transitions, including stomata cell fate transition (Cui et al., 2023), lateral root initiation (Benitez-Alfonso et al., 2013), among others (Bayer and Benitez-Alfonso, 2024). A recent review, mentioned transcription factors involved in microsporogenesis in Arabidopsis, several of which are strong candidates for mobile proteins (Wiese et al., 2024) and should be targeted in future studies.

Callose accumulation has been described as key for PD regulation, thus we evaluated changes in PD frequency in the callose synthase mutant *Osgsl5-3*. We found a reduction in PD frequency at all studied interfaces. *Osgsl5-3* mutant display precocious initiation of premeiotic S-phase and subsequent meiosis aberrations leading to reduced pollen fertility (Somashekar et al. 2023). The finding of reduced PD frequency in *Osgsl5-3* anthers is somewhat surprising but it is plausible that meiotic aberrations in callose-lacking *Osgsl5* anthers are attributable to defects in signalling pathways due to decreased PDs. Our results suggest that callose synthesis and regulation of PDs are required for the control of symplastic signalling to assure timely initiation and normal progression of male meiosis in rice anthers.

### A role for callose biosynthesis in plasmodesmata formation and/or maintenance in rice anthers

Closure or disintegration of PDs has been associated with callose accumulation in neighbouring cell walls (Echlin & Godwin, 1968; Mamun et al. 2005; Steer, 1977; Sager and Lee, 2014), thus we expected more (open) PDs in callose-lacking *Osgsl5-3* mutant anthers. We attempted to study the diffusion of symplast and apoplast fluorescent tracers in anther tissue, however due to technical difficulties symplasmic connectivity is difficult to probe in rice anthers and therefore was not assessed in this study, but PD number was greatly reduced in *Osgsl5-3* anthers at all three cellular interfaces (PMC-PMC, PMC-TC and TC-TC) compared to WT anthers (Fig. 2). The presence of stable PDs on callose-rich PMC walls during meiosis is also documented in male sterile soybeans (Albertsen & Palmer, 1979). Our observation rather supports the hypothesis that callose renders transient stability to PDs during mitosis-meiosis initiation. The properties of callose at PD-associated cell walls has been questioned recently (Abou-Saleh et al. 2018; Amsbury et al., 2018; Bayer and Benitez-Alfonso, 2024), and changes in its role in PD formation likely depend on cell wall composition. We showed that cellulose content remains unaltered in both *Osgsl5-2* and *Osgsl5-3* pre-meiotic and early meiotic prophase I anthers, thus depletion of callose might correlate with a hydrogel model where high cellulose:callose ratio increases cell wall stiffness (Abou-Saleh et al. 2018), potentially affecting the formation (by cell wall digestion) of new PDs. We also found an increase in cell-cell distance between PMCs and PMC-TCs in *Osgsl5* anthers (Fig. 3), thus an alternative (more plausible) explanation is that PDs breakdown as the extracellular space enlarges and cells move apart. Supporting this, we found a negative correlation between PD frequency and cell-cell distance in both WT and *Osgsl5* anthers (Fig. 5). It is also possible that the increase of locular cell volumes driven by anther maturation/elongation (Kelliher & Walbot, 2011), might drive PD breakdown and/or increasing apoplast spacing among anther locular cells independent on callose accumulation. More research is required to fully dissect the role of callose synthesis in PD formation or stability, including for example the ultrastructural analysis of mutants in callose degradation.

### Callose and PD impact cell-cell distances and PMC shaping during mitosis-meiosis transition: a working hypothesis

This study also revealed the impact of callose deposition in the apoplast spacing between anther locular cells during meiosis initiation (Fig. 3) and in PMC shaping in rice meiotic anthers (Fig. 6). Increase in the extracellular space in *Osgsl5* anthers will impact molecular transport via the apoplastic pathway (Roschzttardtz et al. 2013). PDs appear to gradually disappear between PMCs, PMCs and TCs during mitosis-meiosis transition (Fig. 2) and are completely abolished at a later stage (Heslop-Harrison & Mackenzie, 1967; Echlin & Godwin, 1968; Horner and Rogers, 1974) (also see Fig. S2). This observation points to a switch from symplastic to apoplastic communication at the PMCs, as previously proposed in *Lilium* and *Arabidopsis* (Clement & Audran, 1995; Roschzttardtz et al. 2013). An excessive increase of apoplast spacing among meiocytes, and between meiocytes and surrounding nurse cells (Fig. 3) might be responsible for both, changes in symplasmic and apoplastic signalling underlying the *Osgsl5* phenotypes.

One of the *Osgsl5* phenotypes observed in this study was an altered PMC shaping in rice premeiotic anthers. In the TEM images at the premeiotic interphase, we could not observe the intermediate states (i.e., curvy-shaping of central PMC corners), between angular and fully spherical appearances (Fig. 6C, see model Fig. 7). This suggests that callose synthesis and deposition might be required for PMC shaping and sphericalization, by changing, for example, the properties of cell walls in a polarized manner (Fig. 7). It is also possible that the reduced PD number in *Osgsl5* might affect the transport of an unknown signalling factor that triggers sphericalization. This signalling factor could be osmotic (turgor pressure) as in other systems (such as during cotton fibers elongation (Hernandez-Hernandez et al. 2020) the accumulation of callose, and presumed PD closure, is shown to prevent leakage of osmolytes out of the cell changing turgor pressure. Failure in PD closure or the increase in extracellular space in the callose synthesis *Osgsl5-3* mutant may delay an elevation of the turgor pressure sufficient for PMC sphericalization, however this hypothesis needs further validation. Based on our results, we propose that the concentration of OsGSL5-dependent callose at the intercellular space of the anther locular center drives the uneven changes in the extracellular spacing leading to the curvy-shaping appearance of a central PMC corner (Fig. 6A&B, see model Fig. 7).

**Figure 7.**
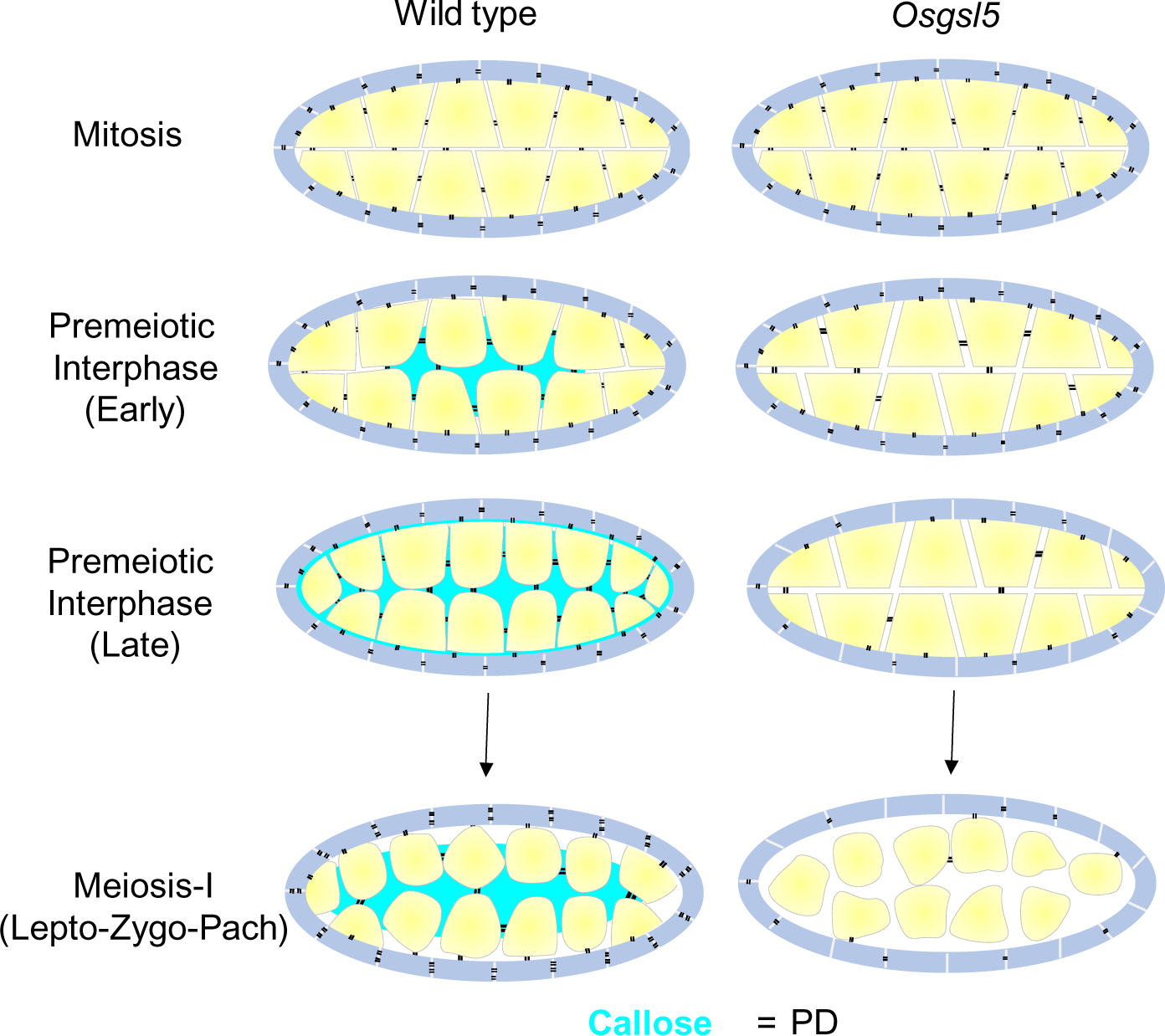
Model representing changes in callose deposition, plasmodesmata connections, cell-cell distance and PMC shaping in anther locules. In wild type (WT), during mitosis and premeiotic interphase stage, PDs are dense among the anther locules cells. Callose deposits at the central locule between angular shaped WT PMCs during premeiotic interphase (Early) and the PMC shape become curvy at the regions corresponding to the callose deposition sites. At premeiotic interphase (Late) stage, callose completely fills the anther locules leading to PMCs sphericalization at meiosis–I. In turn PD number decreases with the increase in callose between PMC-PMC as they reach early meiosis–I. In callose deficient mutant *Osgsl5*, cell-cell distance (apoplastic spaces) increases (but remained constant along the interface) at all premeiotic and early meiosis stages and PD frequency is reduced. The lack of callose is proposed to influence the curvy-shaped PMC corners at premeiotic interphase locule center, delaying sphericalization affecting meiosis.

Taken together, this study sheds light on the biological significance of the cellulose-to-callose switch in the walls of the PMCs in premeiotic anther locules and the profound impact of callose on the number of PD connections, apoplast spacing and germ cell wall shaping during mitosis-meiosis transition (Fig. 7). Our results link PD frequency controlled by callose deposition with successful meiosis and plant reproduction and further raises several interesting research questions for future studies.

## Acknowledgements

We thank Tsuyoshi Koide Ph.D. (Mouse Genomics Resources Laboratory, National Institute of Genetics) for coordinating the electron microscopic collaboration. We thank Misako Kasihara for helping with the electron microscopic experiments.

## Author contributions

HS and KIN conceptualized the work. HS, KT, AO, RH performed the experiments, HS, KT, YBA and KIN designed experiments and analysed the data, HS, YBA, KIN contributed in the writing-reviewing and editing the manuscript.

## Data availability

All data supporting this study are available upon reasonable request to the corresponding authors.

